# Differential Expression and Interaction network construction of Noncoding RNA and mRNA in Heart Tissue associated with Tetralogy of Fallot

**DOI:** 10.1101/2020.02.18.953877

**Authors:** Hongdan Wang, Cunying Cui, Yanan Li, Yuanyuan Liu, Taibing Fan, Bangtian Peng, Lin Liu

**Author notes:** Corresponding authour: Lin Liu.

## Abstract

Tetralogy of Fallot (TOF) is still the most common and complicated cyanotic congenital heart defect of all congenital heart diseases with a 10% incidence. Surgery repair is often necessary in infancy. The etiology of TOF is complex and genetic and epigenetic mechanisms such as chromosomal abnormalities, gene mutations, nucleic acid modifications, non-coding RNA, and circular RNA(circRNA) play an important role in its occurrence. RNA not only plays an auxiliary role of genetic information carrier, but also plays a more important role in various regulatory functions. There are few studies on the action mechanism of non-coding RNA. Aim to gain more in-depth knowledge of TOF, we collected tissue samples of the right ventricular outflow tract of 5 TOF children with no other intracardiac and extracardiac malformations and 5 normal fetuses. We systematically analyzed the specific long non-coding RNA (lncRNA), microRNA(miRNA), circRNA and messenger RNA(mRNA) profiles of TOF. To our knowledge, there are no reports of genome-wide study of transcriptome in TOF and we first obtained meaningful differentially expressed lncRNAs, miRNAs, circle RNA and mRNAs.

---

The total incidence of neonatal defects in China is 5.6%, and 0.9 million neonates are born with defects every year. The incidence of neonatal defects is increasing annually. The heart is the first major internal organ in the process of embryogenesis and is important for the survival of an embryo. Congenital heart disease (CHD) has ranked first for five consecutive years in neonates. The overall incidence of CHD in the population is 1‰–13‰. In China, the incidence of CHD is 4‰–8‰. Children with severe CHD die in childhood, thereby causing psychological trauma and economic burden to families and society. Tetralogy of Fallot (TOF) remains the most common complex cyanotic congenital heart defect among all congenital heart diseases given its 10% incidence (Bedard *et al.* 2009). Surgery repair is often necessary in infancy, and the perioperative mortality is less than 5%. However, long-term follow-up has shown that patients who have undergone total repair of TOF are often at high risk for restenosis of the right ventricular outflow tract (RVOT), pulmonary regurgitation, and ventricular dysfunction, and late sudden cardiac death may occur in 0.5% to 6% of patients due to ventricular tachycardia or other unknown causes(Massin *et al.* 2011; Zhang *et al.* 2013; Latus *et al.* 2015; Shin *et al.* 2016). The exact etiology of TOF remains unclear, and its pathogenesis is complex. Genetic and epigenetic mechanisms, such as chromosomal abnormality, gene mutation, nucleic acid modification, noncoding RNA (ncRNA), and circular RNA (circRNA), play an important role in its occurrence. At present, chromosomal abnormalities, gene mutations, and other genetic mechanisms have been deeply studied and widely used in the diagnosis and treatment of clinical diseases. However, the action mechanism of ncRNAs on TOF needs to be further studied.

Recently, genome-wide transcriptional studies have represented that only almost 1% of the human genome produces biologically meaningful RNA transcripts, while a much larger proportion of the genome is transcribed into ncRNAs(Ponting *et al.* 2009). RNA not only plays an auxiliary role as a carrier of genetic information but also demonstrates various regulatory functions. In comparison with the diversity of RNA transcripts which encode for proteins, the functions of a small number of ncRNAs have been demonstrated. ncRNAs refer to RNA that is not translated into protein. ncRNAs involved in gene regulation include miRNA, long-chain ncRNA (lncRNA), circRNA, PIWI interacting RNA, and small interfering RNA(Bajpai *et al.* 2010). Among them, lncRNAs, microRNAs (miRNAs), and circRNAs have attracted the most attention.

lncRNAs can competitively bind miRNA through competitive endogenous RNA (ceRNA) to relieve the translation inhibition of miRNA on the target gene messenger RNA (mRNA) (Salmena *et al.* 2011). The expression of lncRNAs are not only cell type and tissue-specific. Some are expressed only at specific stages of eukaryotic development. Sufficient evidence has shown that lncRNAs are key molecules that regulate gene expression, chromatin number maintenance, transcriptional regulation, and post regulation (Tano and Akimitsu 2012; LI *et al.* 2013). lncRNAs play significant parts in various organisms’ physiological and pathological processes, such as embryonic development, organogenesis, and tumorigenesis (Jariwala and Sarkar 2016). In recent years, increasing attention has been paid to research on lncRNAs in the cardiovascular field. Studies have demonstrated that lncRNAs play a crucial part in promoting cardiomyocyte differentiation and maintaining cardiac function (Schonrock *et al.* 2012; Philippen *et al.* 2015; Gao *et al.* 2017).

miRNAs are a kind of endogenous noncoding, single-chain, and small molecular RNA composed of approximately 22 nucleotides. They widely exist in eukaryotic cells and are highly conserved. miRNAs are a multifunctional regulator of gene expression in higher eukaryotes. The regulation mode is that the targeted binding of mRNA is mainly conducted, thereby further inhibiting or blocking translation(Yates *et al.* 2013). Heart development is a process of polygenic regulation. Studies have shown that miRNAs can participate in the expression, transcription, splicing, and translation of important genes in the development of the heart and in the regulation of related pathways, which may lead to heart defects (O’brien *et al.* 2012).

circRNA is a kind of special RNA molecule. Different from traditional linear RNA, circRNA has a closed ring structure, which is not affected by RNA exonuclease and is difficult to degrade. Its expression is more stable compared with traditional linear RNA. New evidence suggests that circRNAs are widely expressed in mammalian cells and exhibit cell or tissue specificity. circRNAs have been proven to get involved in regulating various biological processes(Yu and Kuo 2019). circRNAs can also affect cell physiological processes through various molecular mechanisms. They can be used as mediator of miRNA and RNA binding protein to regulate protein expression and translation. Similar to lncRNAs, circRNAs are rich in miRNA binding sites. They can play the role of miRNA through mechanism competition of ceRNA and relieve miRNA inhibition on their target gene. As such, the target gene’s expression level can be increased. circRNAs play a significant regulatory part in diseases through their interaction with miRNAs associated with diseases or other functional elements (Han *et al.* 2018). However, our understanding of the regulation and function of circRNAs is limited.

TOF is the most common cyanotic congenital heart disease. Most of TOF have nothing to do with chromosome or microduplication/deletion. The pathogenesis of simple TOF has not been elucidated. However, ncRNAs may play a significant regulatory part in heart formation. Little is known regarding the changes in the transcriptome and the possible associations with TOF. Thus, we systematically analyzed the specific lncRNA, miRNA, circRNA, and mRNA profiles of TOF to gain more in-depth knowledge. As we know, no genome-wide study has concentrated on the transcriptome in TOF. This is the first work to obtain meaningful differentially expressed lncRNAs, miRNAs, circle RNA, and mRNAs.

## Materials and methods

### Subjects

The case group included five children with TOF (two boys and three girls, aged 6–22 months). Cyanosis was found in all children, and rough systolic murmur was found between the second and fourth intercostals at the left border of the sternum by auscultation. Echocardiography showed ventricular septal defect, aortic straddle, RVOT obstruction, pulmonary artery stenosis, right ventricular enlargement, and right ventricular anterior wall thickening. The children and their parents had no other intracardiac and extracardiac malformations. In the control group, five fetuses, including three males and two females with gestational age of 25–28 weeks, had normal hearts. Given fetal distress, cord twist, and placental abruption, odinopoeia was conducted.

### Extraction of RNA

The Total RNA including small RNA was collected from 10–20 mg of frozen tissue of the right ventricle by using a TRIzol reagent (Invitrogen, Gaithersburg, MD, USA), which was purified with a mirVana miRNA Isolation Kit (Ambion, Austin, TX, USA) according to the instructions of the manufacturer. RNA’s purity and concentration were defined by OD260/280 by using a spectrophotometer (NanoDrop ND-1000). RNA integrity was defined by 1% formaldehyde denaturing gel electrophoresis. RNA integrity was defined by capillary electrophoresis with an RNA 6000 Nano Lab-on-a-Chip kit and Bioanalyzer 2100 (Agilent Technologies, Santa Clara, CA, USA). Only if the integrity number values of RNA were more than 6, it was analyzed later.

### Microarray

The specific lncRNA, mRNA, miRNA, and circle RNA profiles of TOF were analyzed by using Agilent human lncRNA+mRNA Array v4.0, Agilent human miRNA Array V21.0, and Agilent human circRNA Array v2.0.

lncRNA and mRNA profiling was performed using the Agilent human lncRNA+mRNA Array V4.0 designed with four identical arrays per slide(4×180k), which included the number of probes of human lncRNAs and human mRNAs being approximately 34,000 and 41,000 respectively. lncRNA and mRNA target sequences were selected from diverse databases such as GENCODE/ENSEMBL, Human LincRNA Catalog [1], RefSeq, USCS, NRED (ncRNA Expression Database), LNCipedia, H-InvDB, LncRNAs-a (Enhancer-like), Antisense ncRNA pipeline, etc.. Repeated probes were used to detect each RNA for two times. The control probes were contained in the array.

The Agilent miRNA array (8 × 60K) was used to detect miRNA profiling, which included 2549 probes of human mature miRNAs and the miRNA target sequences were from the miRBase R21.0. Repeated probes were used to detect each miRNA for times. The control probes were contained in the array.

The Agilent human circRNA Array V2.0 (4 × 180K) was used to detect circRNA profiling, which included approximately 170,340 probes of human circRNAs and the circRNA target sequences were from circBase and DeepBase. Long and short probes were used to detect each circRNA at the same time. The control probes were contained in the array.

### RNA amplification, labeling, and hybridization

#### Agilent human lncRNA+mRNA Array v4.0

Cy5 and Cy3-dCTP was used to label the cDNA) by using the Eberwine’s linear RNA amplification method and subsequent enzymatic reaction (Patterson *et al.* 2006). By using the CbcScript reverse transcriptase with cDNA synthesis system according to the manufacturer’s protocol with T7 Oligo (dT) and T7 Oligo (dN), double-stranded cDNAs were synthesized using 1 μg of total RNA.

DNA polymerase and RNase H were used for double-stranded cDNA (dsDNA) synthesis PCR NucleoSpin Extract II Kit eluted with 30 μL elution buffer was used for the purifying of the dsDNA products. The products were concentrate to 16 μL and conducted with vitro transcription reactions in 40 μL using the T7 Enzyme Mix at 37 °C for 14 h. the RNA Clean-up Kit was used for purifying the amplified cRNA.

Klenow enzyme labeling was performed next by using CbcScript II reverse transcriptase. In short, the amplified RNA (2 μg) was mixed with random nanomer (4 μg), denatured for 5 min at 65 °C, and immediately cooled on the ice. Then, 4× first-strand buffer (5 μL), o0.1M DTT (2 μL), and CbcScript II reverse transcriptase (1.5 μL) were mixed and added. The mixtures were incubated for 10 min at 25 °C ad then for 90 min at 37 °C. The purification method of cDNA products was as follows. cDNA was added with random nanomer (4 μg), 95 °C for 3 min, and cooled on ice immediately for 5 min. Then, Klenow buffer (5 μL), dNTP, and Cy5-dCTP or Cy3-dCTPwere mixed and added. Then, added Klenow enzyme for1.2 μL, and incubated for 90 min at 37 °C. Labeled cDNA was purifiedas follows. The hybridization solution of the labeled controls samples and test samples were dissolved in 80 μL including 3× SSC, 0.2% SDS, 5× Denhardt’s solution, and 25% formamide. DNA in the hybridization solution was denatured at for 3 min 95 °C before loading on the microarray. Arrays were hybridized overnight at 42 °C in an Agilent Hybridization Oven at the rotation speed of 20 rpm. At last, the microarray was washed using the consecutive solutions (0.2% SDS, 2× SSC and 0.2× SSC).

#### Agilent human miRNA Array V21.0

The experiments were carried out in accordance with the protocol of the manufacturer. In short, the miRNA labeling reagent was used to label the miRNAs. pCp-Cy3 was used to dephosphorylate and ligate 200 ng total RNA. Next, we purify and hybridize the labeled RNA. Last, the Agilent microarray scanner was used to scan the images and Agilent feature extraction software version 10.10 was used to grid and analyze them.

#### Agilent human circRNA Array v2.0

Total RNAs were first digested by using Ribonuclease R to ensure the specific and efficient labeling of cirRNA. Then, Cy3-dCTP was used to label cDNA in accordance with Eberwine’s linear RNA amplification method. The succeeding enzymatic reaction was as described above. The next steps of RNA amplification, labeling, and hybridization are described in the “Agilent human lncRNA+mRNA Array v4.0” section.

### Microarray imaging and sample analysis

The Agilent GeneSpring software V13.0 was used to summary, normalize and control in quality the data of the lncRNA+mRNA array, miRNA array, and circRNA array.

The genes which were differentially expressed were selected by using the Threshold values of ≥2 and ≤-2-fold change and a corrected P value of Benjamini–Hochberg, less than 0.05, which had significant differences. Log2 was used to process the data and the CLUSTER 3.0 software was used to adjust the data. The hierarchical clustering was used to analyze the data and cluster heat maps were generated. At last, the Java TreeView (Stanford University School of Medicine, Stanford, CA, USA) was used to conduct the tree visualization. On the basis of the normalized data, the Pearson correlation coefficient matrix of the correlation between samples was made. The box plot was drawn, showing the gene expression of different samples before and after data normalization. Principal component analysis (PCA) was used to reflect the similarity of samples, and the expression of the samples was shown in the 3D space through the dimension reduction of data.

### Comparative analysis of case and control groups

T-test was used to analyze the difference between groups. The default screening criteria for significant difference in this study were as follows: when the gene was upregulated, the number of detected samples required to account for more than 60% of the total number of this group. When the gene was downregulated, the requirement was the same. The difference multiple FC(ABS)≥2.0 and *P*≤0.05 were also required. The larger the difference between the multiple of FC (ABS), the greater the difference between the two groups. The lower the *P* value, the higher the reliability of differential genes. Cluster analysis was performed, and a scatter map and volcano map were drawn to illustrate the differences between groups.

### Construction of the coding–noncoding gene co-expression (CNC) network

#### Construction of the lncRNA–mRNA network

The correlation analysis between the lncRNA and mRNA differentially expressed was used for constructing the network of CNC. For construction of the network, Pearson correlation was calculated, and the critical correlation lncRNA and mRNA pairs were chosen. When Pearson correlation coefficients, which are not less than 0.99, the lncRNAs and mRNAs pairswere selected to draw the network by using software Cytoscape. In the network construction analysis, a degree was the simplest and most important measure of a gene centrality which was defined as the number of connections between one node **to** the other(Barabasi AND OLTVAI 2004).

The lncRNA prediction for Cis-acting and trans-acting was performed in this study. The prediction of Cis-acting lncRNA was conducted by their correlation to the expressed protein-coding genes. When the Pearson’s correlation coefficient less than 0.99, Cis-acting lncRNA could be selected. The lncRNA resides at the genomic loci where a protein-coding gene and an lncRNA gene are within 10 kb of each other along the genome(Jia *et al.* 2010).

Trans-prediction was performed by using blat tools (http://hgdownload.cse.ucsc.edu/admin/exe/) with the setting of the default parameter to compare the lncRNA with its co-expression mRNAs’ 3′ UTR.,

#### Construction of the miRNA–mRNA network

miRNA target gene prediction was analyzed by using miRWalk2.0, which gave the prediction results of 12 miRNA target gene prediction programs. The programs included miRWalk, DIANA-microTv4.0, miRanda-rel2010, mirBridge, miRDB4.0, miRmap, miRNAMap, PicTar2, PITA, RNA22v2, RNAhybrid2.1, and Targetscan 6.2.

#### Construction of the circRNA–miRNA network

CircRNAs play crucial parts in the function and transcriptional control of miRNA through acting as ceRNAs or positive regulators on their parent coding genes. The network of circRNA–miRNA was performed based on the miRanda-3.3 software. The open-source bioinformatics software Cytoscape was used for constructing the network.

### Gene annotation and enrichment analysis

Gene Ontology (GO), Kyoto Encyclopedia of Genes and Genomes (KEGG) pathway enrichment, and disease analyses were conducted for the target gene mRNA of significantly different lncRNAs, miRNAs, and circRNAs. In the KEGG pathway enrichment and disease analyses, the first 20 critically enriched GO terms were selected, and the first 20 significantly enriched terms from biological process (BP), molecular function (MF), and cellular component (CC) were selected in GO. The histogram drawn on the basis of the P value could directly reflect significantly enriched terms. A bubble chart was also provided to show the enrichment analysis results. Finally, candidate signal pathways related to TOF were used to further determine candidate lncRNAs, mRNAs, miRNAs, and circRNAs.

### Quantitative real-time polymerase chain reaction (qRT-PCR)

The RNA was extracted by using the TRIzol method, which was reverse, transcribed to cDNA by using a TaKaRa reverse transcription kit. Reaction amplification was performed by using ABI StepOne Real-Time PCR system (ABI, Vernon, USA). The reaction conditions were as follows: 50 °C, 2 min; 95 °C, 10 min; 95 °C, 20 s; 60 °C, 30 s; and 72 °C, 30 s for a total of 40 cycles; 95°C, 15 s, 60°C, 1 min, 95°C, 15 s.

### Data Availability Statement

The datasets analyzed for this study can be found in a shared Microsoft OneDrive network disk (https://1drv.ms/f/s!Akmkg4ZxWXm5jbJis0B97A2RBPPr-g)

## Results

### lncRNA/mRNA

According to the screening criteria mentioned previously (FC≥Ccordin0.05), altogether of 3228 lncRNAs which are differentially expressed and 4295 mRNAs which are differentially expressed were found. Among them, 1683 were upregulated lncRNAs, and 1545 were downregulated lncRNAs; by contrast, 2817 mRNAs were upregulated, and 1478 mRNAs were downregulated (Attached Lists 1 and 2). The clustering analysis result (Fig. 1) suggests that in-group samples have a good correlation and high consistency, whereas the between-group samples are significantly different and can be clearly distinguished. Thus, the selection feasibility of tissue samples is high. Co-expression analysis was conducted on the selected lncRNAs that are differentially expressed and mRNAs. A coexpression network map was constructed (Fig. 2), and 44 lncRNAs and 497 mRNAs co-expressed were identified. The target genes of the co-expressed lncRNA and mRNA were predicted, and the coexpression network map was constructed (Fig. 3). Finally, 66 lncRNAs and 62 mRNAs that may target each other were identified.

**Figure 1.**
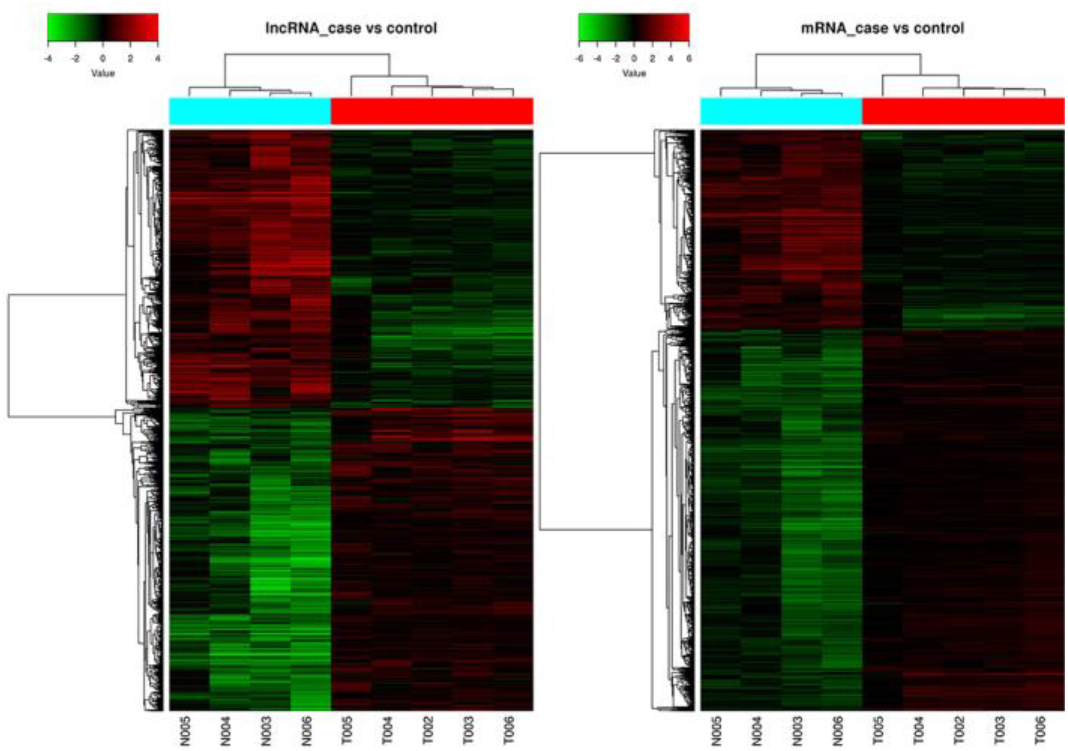
The sample clustering analysis results of lncRNAs and mRNAs

**Figure 2.**
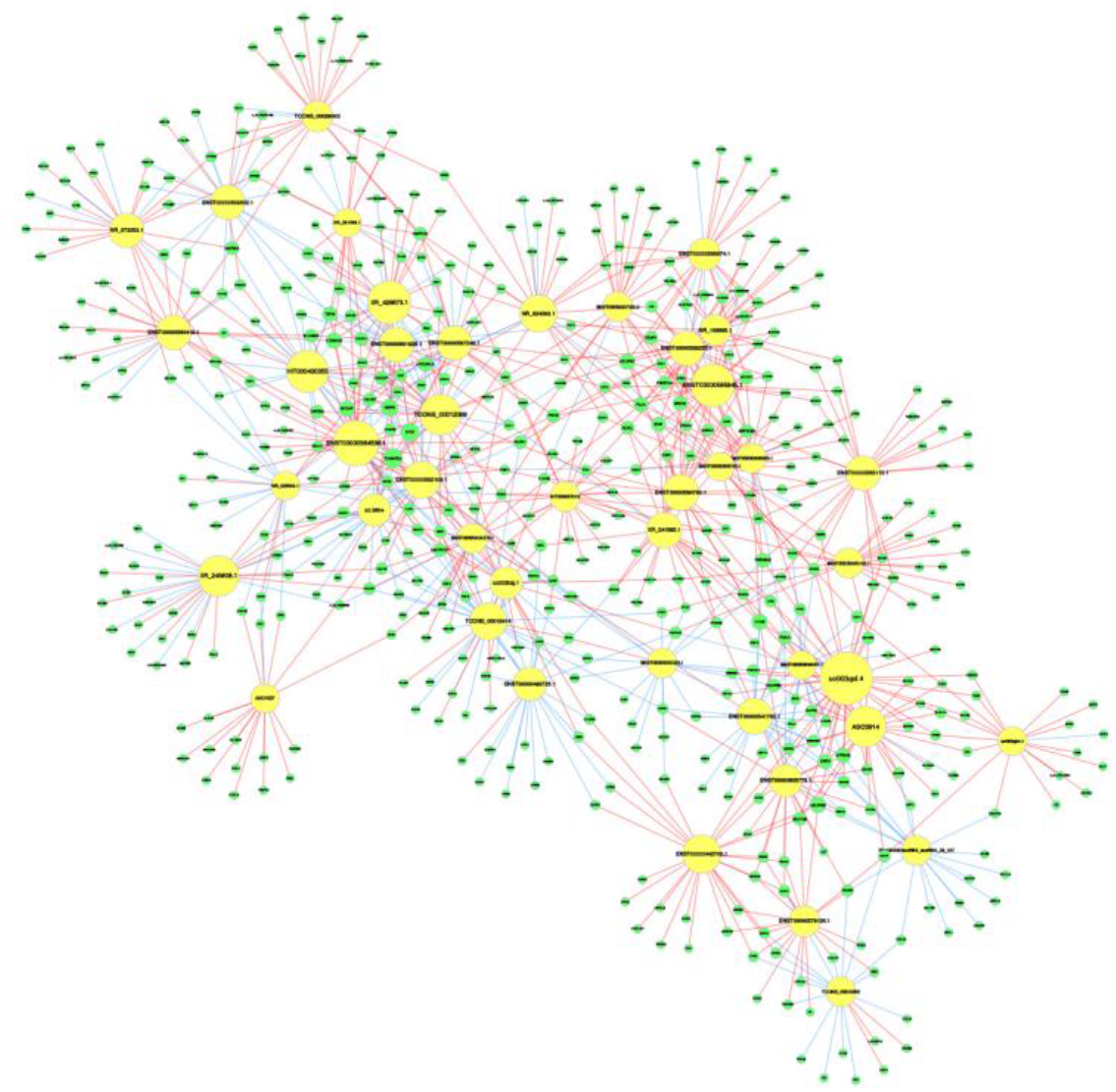
The co-expression network map of lncRNAs and mRNAs

**Figure 3.**
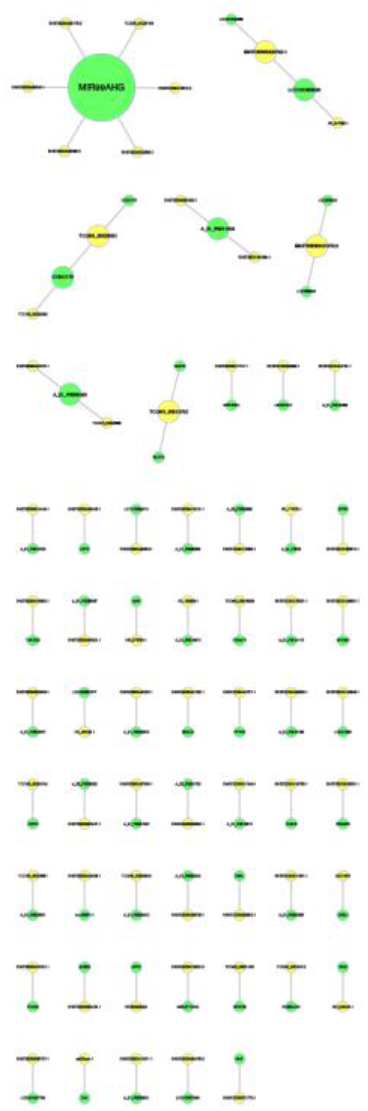
The target genes of the co-expressed lncRNA and mRNA

GO, KEGG pathway enrichment, and disease analyses were conducted for the candidate genes that target each other. In GO analysis, 20 significantly enriched terms were selected from BP, MF, and CC. A histogram was drawn on the basis of the P value, as shown in Fig. 4. In KEGG pathway enrichment and disease analyses, a histogram was drawn for the top 20 terms, as shown in Fig. 5. A total of five lncRNAs and mRNAs that interacted with one another were selected finally (Table 1).

**Figure 4.**
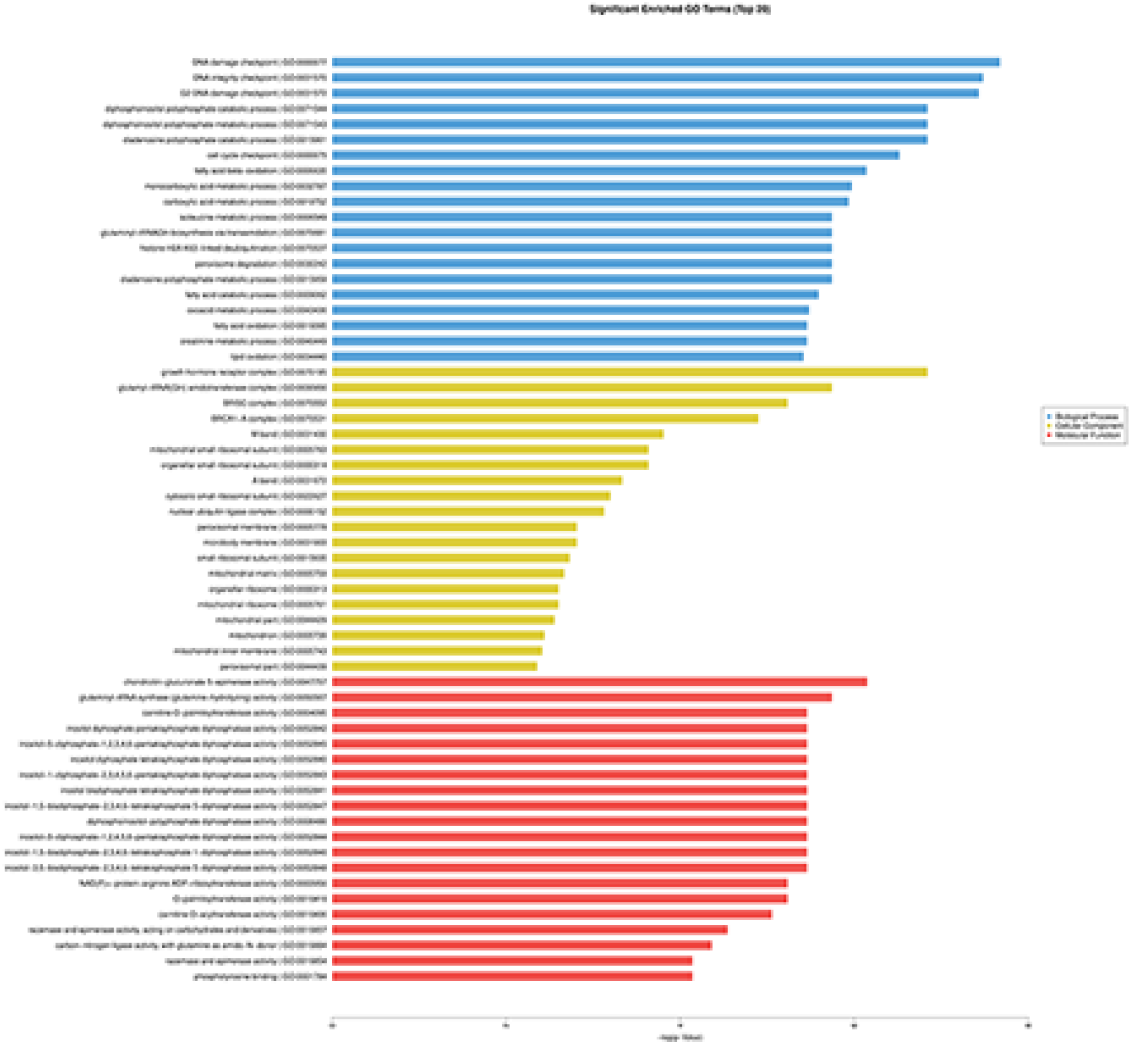
A histogram of 20 significantly enriched terms selected from BP, MF, and CC on the basis of the GO analysis

**Figure 5.**
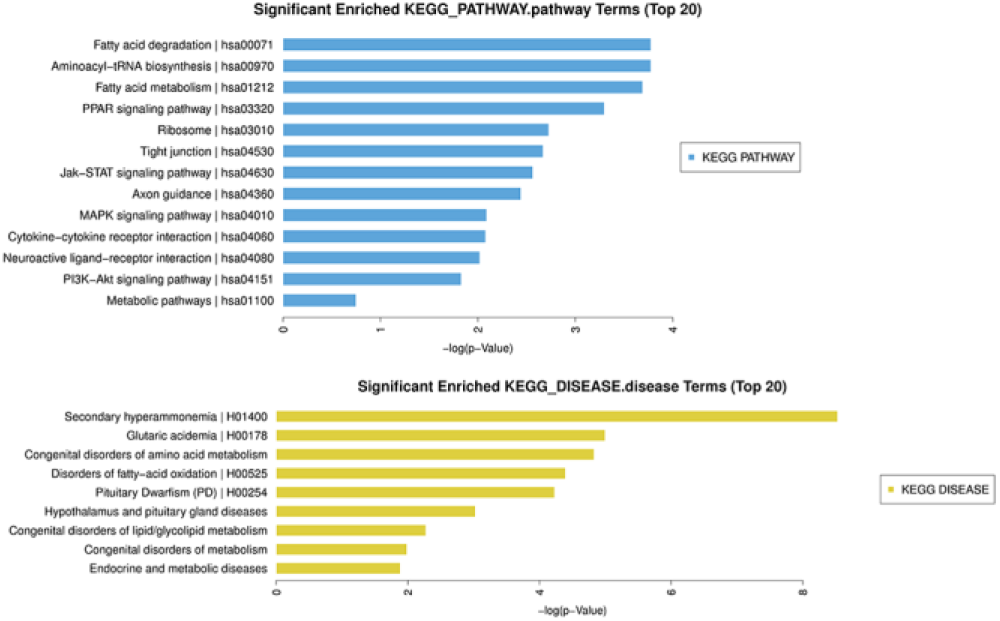
A histogram of top 20 terms by KEGG pathway enrichment and disease analyses

**Table 1.** Five selected lncRNAs and their mRNA that may be the target genes.

| LncRNA ID | LncRNA Gene name | target mRNA Gene<br>name | mRNA RefSeq<br>Accession | Case vs<br>Control ( lncRNA ) | Case vs<br>Control ( mRNA ) |
| --- | --- | --- | --- | --- | --- |
| ASO1873 | ASO1873 | QRSL1 | NM_018292 | up | up |
| ENST00000452466.1 | ENSG00000236723.1 | CPT2 | NM_000098 | up | up |
| NR_073058.1 | C7orf55 | GHR | NM_000163 | up | up |
| ENST00000517747.1 | ENSG00000245281.2 | MRPS18C | NM_016067 | up | up |
| uc002qim.1 | AL833150 | ZAK | NM_133646 | up | up |

### miRNA

A total of 118 differentially expressed miRNAs, including 94 upregulated and 26 downregulated miRNAs, were screened (Attached List 3). A Circos plot was drawn on the basis of the differences (Fig. 6). This plot shows the degree of difference among genes on the basis of their location. Figure 6 shows 37 upregulated miRNAs and 10 downregulated miRNAs with a high degree of difference. The number 13, 15, and Y chromosomes are not distributed. Clustering analysis and principal component analysis suggest that in-group samples have a good correlation and a high consistency. The between-group samples are significantly different and can be clearly grouped, with strong data reliability. GO, KEGG pathway enrichment, and disease analyses were conducted for different genes (Figs. 7 and 8). Target gene prediction was conducted for differential miRNA. miRNAs that interacted with another and conformed to the expression trend of the target mRNA were selected (Table 2).

**Figure 6.**
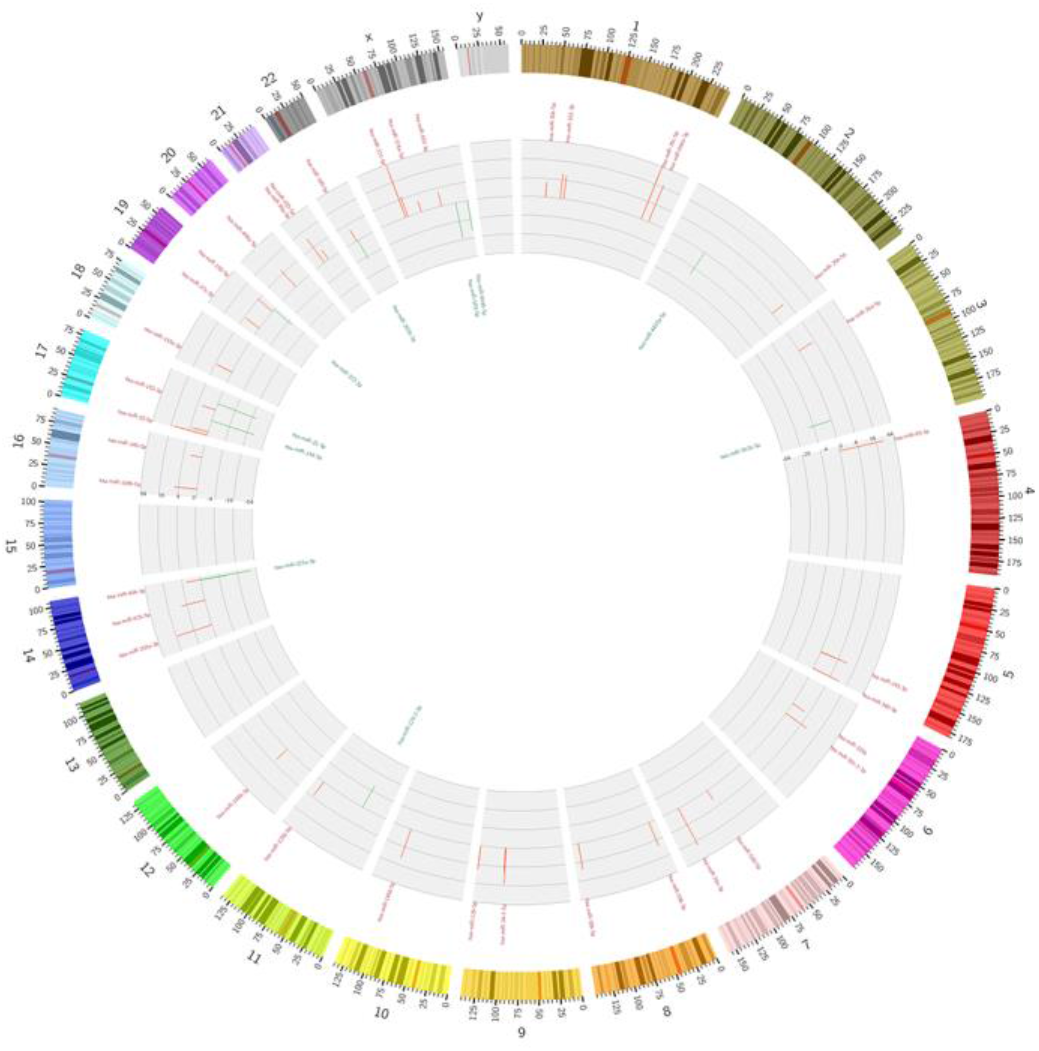
A Circos plot shows 37 upregulated miRNAs and 10 downregulated miRNAs with a high degree of difference on the basis of their location

**Figure 7.**
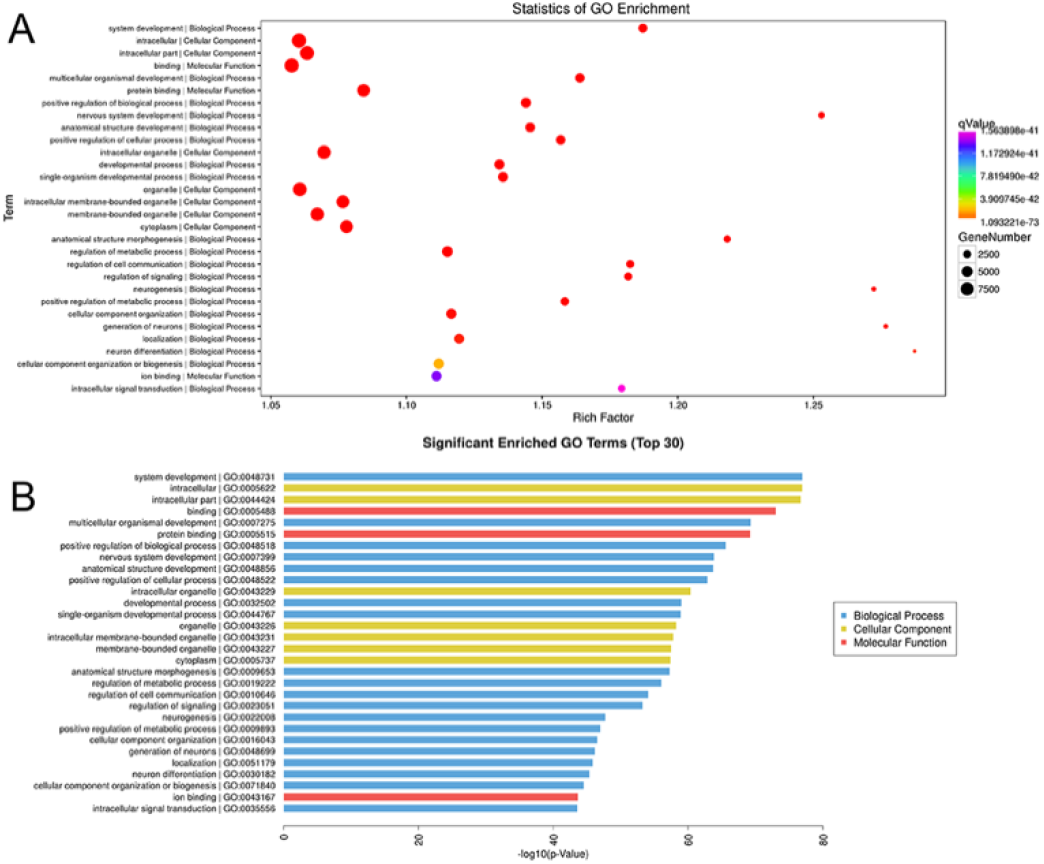
Statistics of GO Enrichment and significant enrichment Go terms of TOP 30 differentially expressed miRNA

**Figure 8.**
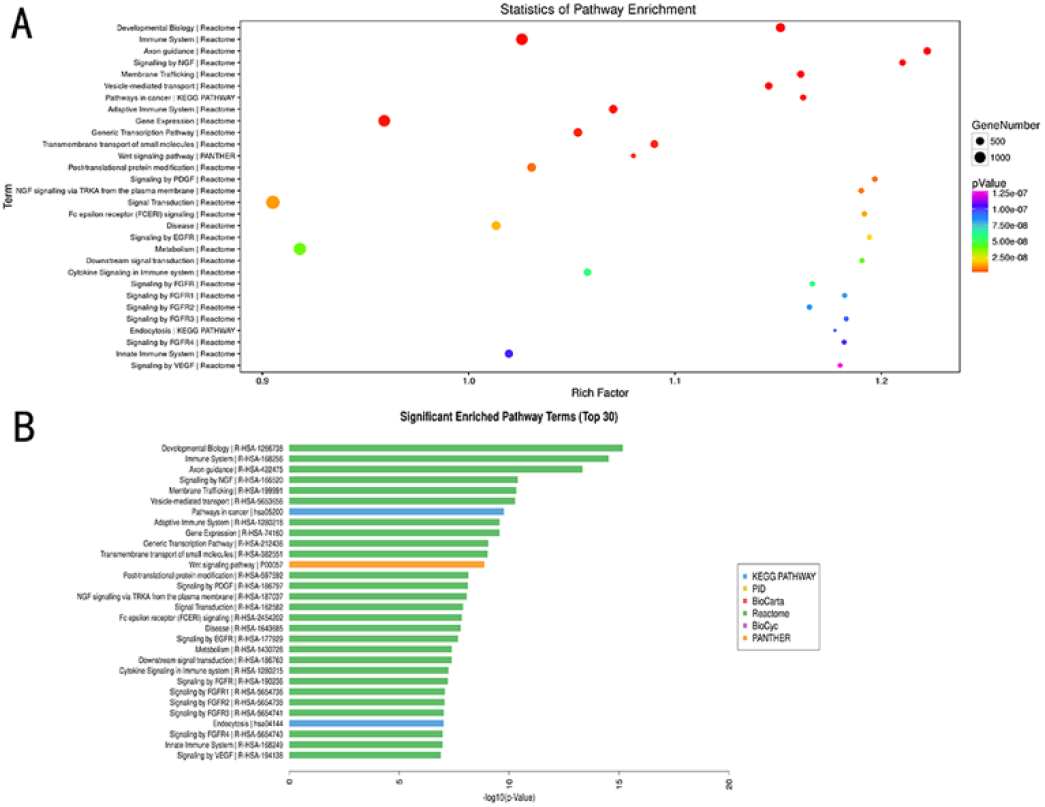
Statistics of KEGG Pathway Enrichment and significant enrichment KEGG Pathway terms of TOP 30 differentially expressed miRNA

**Table 2.**
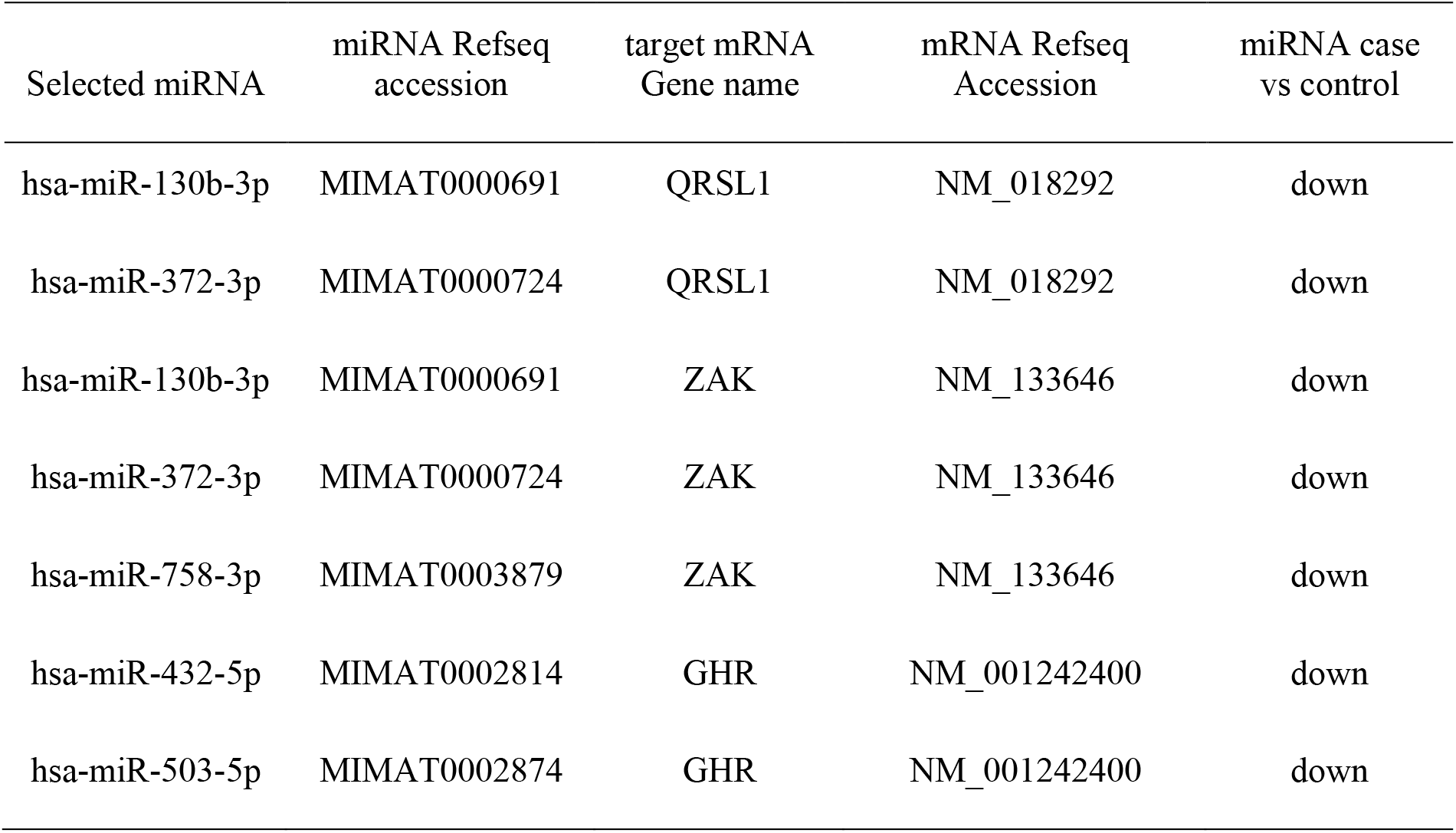
Seven selected miRNAs and their mRNA that may be the target genes.

| Selected miRNA | miRNA Refseq<br>accession | target mRNA<br>Gene name | mRNA Refseq<br>Accession | miRNA case<br>vs control |
| --- | --- | --- | --- | --- |
| hsa-miR-130b-3p | MIMAT0000691 | QRSL1 | NM_018292 | down |
| hsa-miR-372-3p | MIMAT0000724 | QRSL1 | NM_018292 | down |
| hsa-miR-130b-3p | MIMAT0000691 | ZAK | NM_133646 | down |
| hsa-miR-372-3p | MIMAT0000724 | ZAK | NM_133646 | down |
| hsa-miR-758-3p | MIMAT0003879 | ZAK | NM_133646 | down |
| hsa-miR-432-5p | MIMAT0002814 | GHR | NM_001242400 | down |
| hsa-miR-503-5p | MIMAT0002874 | GHR | NM_001242400 | down |

### circRNA

A total of 18,016 circRNAs which are differentially expressed, including 9404 upregulated circRNAs and 8612 downregulated circRNAs, were screened. On the basis of differences, a Circos plot was drawn (Fig. 9). This plot is based on the location of genes. It can show the degree of difference between differential genes. The plot was distributed in all chromosomes. Clustering analysis and PCA were conducted for cases and control samples. The results show that in-group samples have a good correlation and high consistency; between-group samples are significantly different and can be clearly grouped, and data have strong reliability. GO, KEGG pathway enrichment, and disease analyses were conducted for differential circRNAs (Figs. 10 and 11). The most prominent function of circRNAs is its role as miRNA sponge to regulate target gene expression by inhibiting miRNA activity. A circRNA–miRNA network was constructed (Fig. 12). The figure shows the interaction network between the first eight circRNAs and multiple target miRNAs. In these eight circRNAs, except for hsa_circ_0056861_CBC1, the expression of the remaining circRNAs was downregulated compared with that in the control group. Three differentially expressed circRNAs, namely, hsa-circRNA301, hsa_circ_0113626, and hsa_circ_0129079, may be related with the target mRNA and were selected. The target mRNA of hsa-circRNA301 and hsa_circ_0113626 is CPT2, and that of hsa_circ_0129079 is GHR.

**Figure 9.**
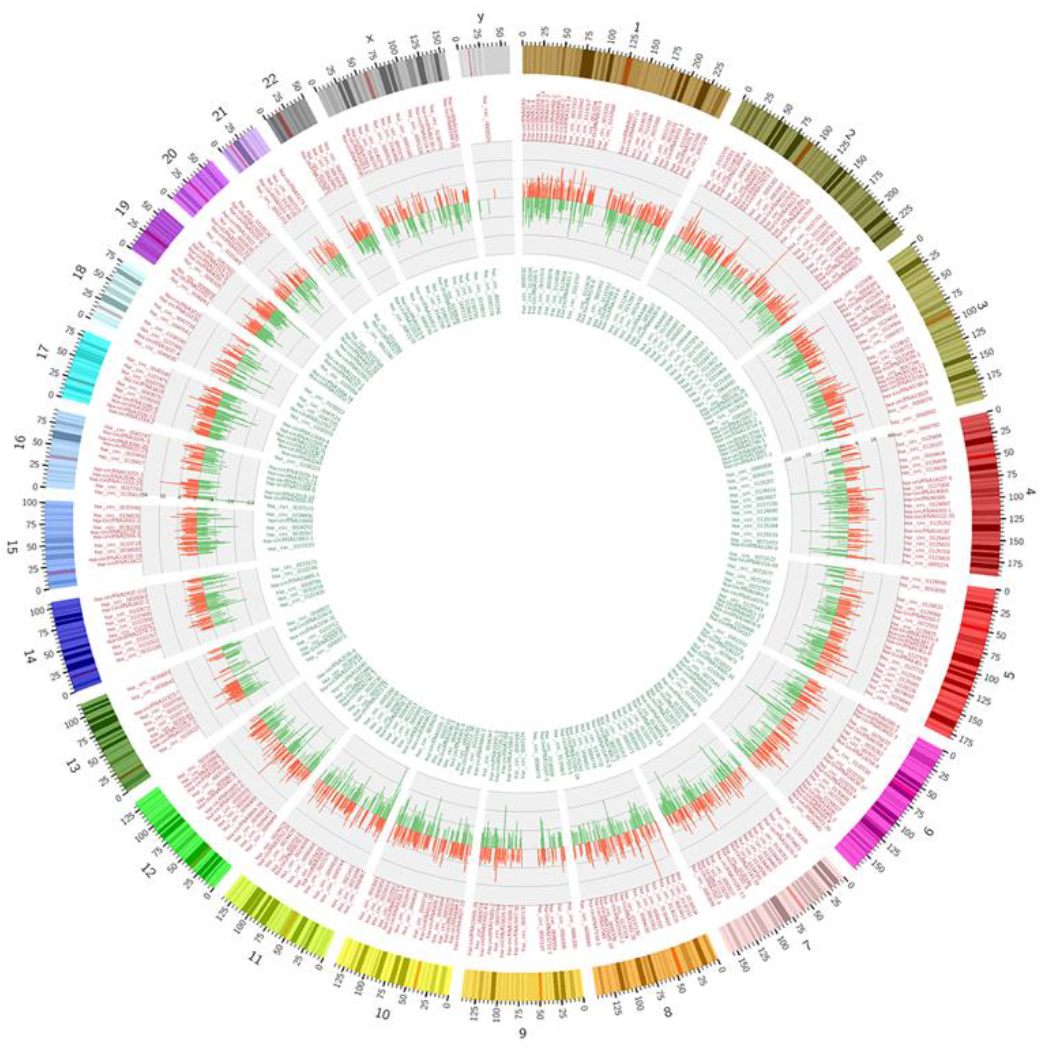
A Circos plot shows differentially expressed circRNAs on the basis of their location

**Figure 10.**
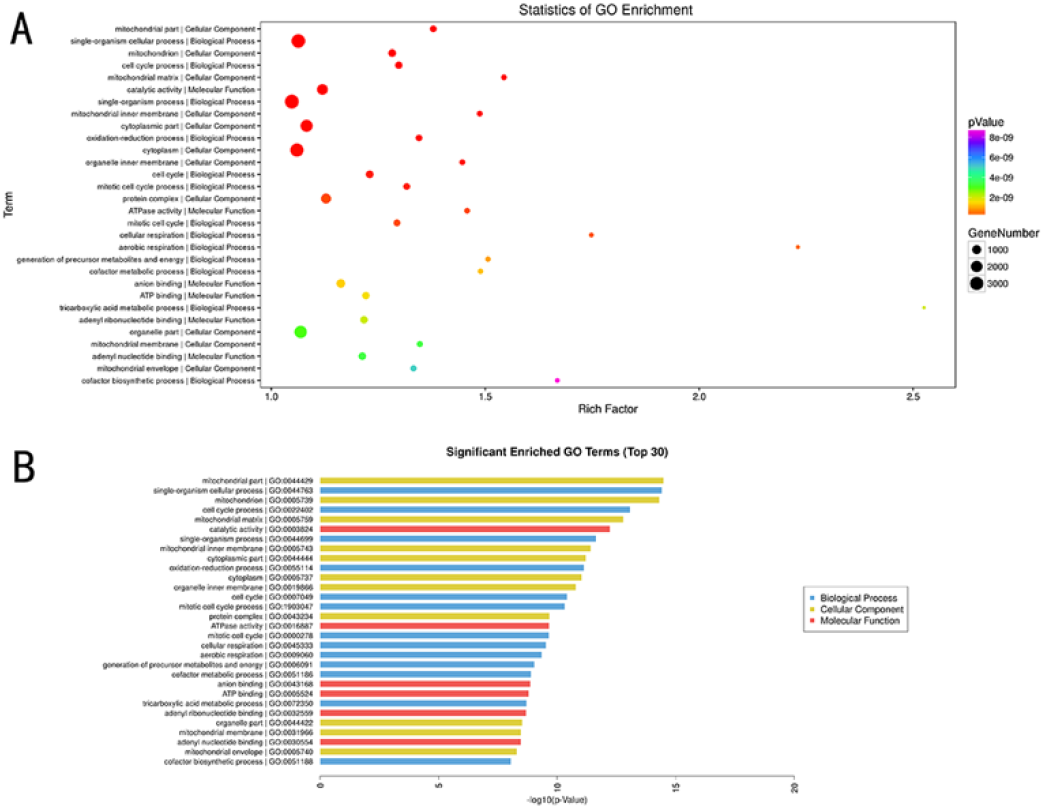
Statistics of GO Enrichment and significant enrichment Go terms of TOP 30 differentially expressed circRNAs

**Figure 11.**
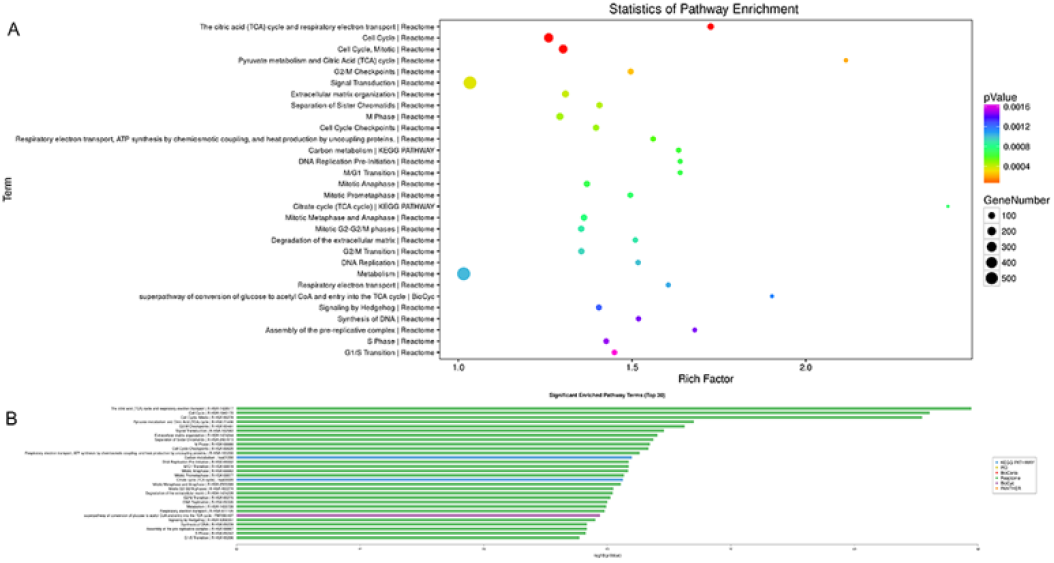
Statistics of KEGG Pathway Enrichment and significant enrichment KEGG Pathway terms of TOP 30 differentially expressed circRNAs

**Figure 12.**
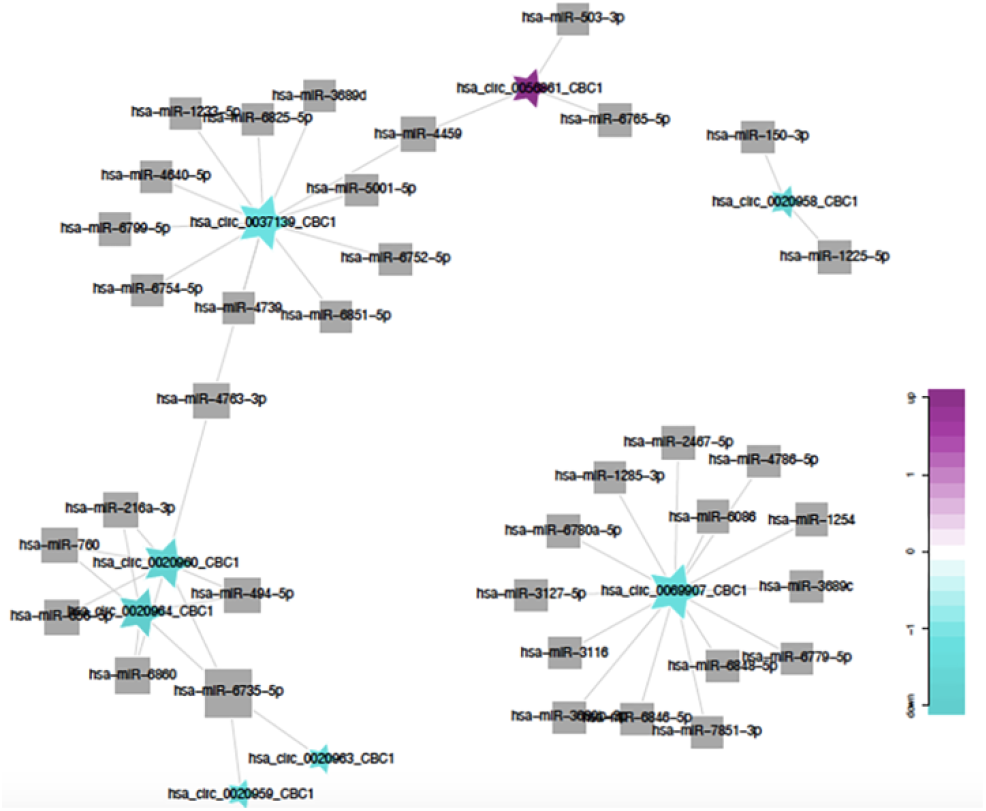
The circRNA–miRNA network shows the interaction network between the first eight circRNAs and multiple target miRNAs

### qRT-PCR

qRT-PCR was conducted for the five pairs of candidate lncRNA/mRNAs in the case and control groups. The test result shows that the relative quantitative analysis results of four lncRNAs and four mRNAs are consistent with the chip results (Fig. 13). The four differential candidate lncRNAs are ASO1873, ENST00000452466.1, NR_073058.1, and uc002qim.1, except for ENST00000517747.1. The four differential mRNAs include QRSL1 (NM_018292), CPT2 (NM_000098), GHR (NM_000163), and ZAK (NM_133646), except for MRPS18C (NM_016067).

**Figure 13.**
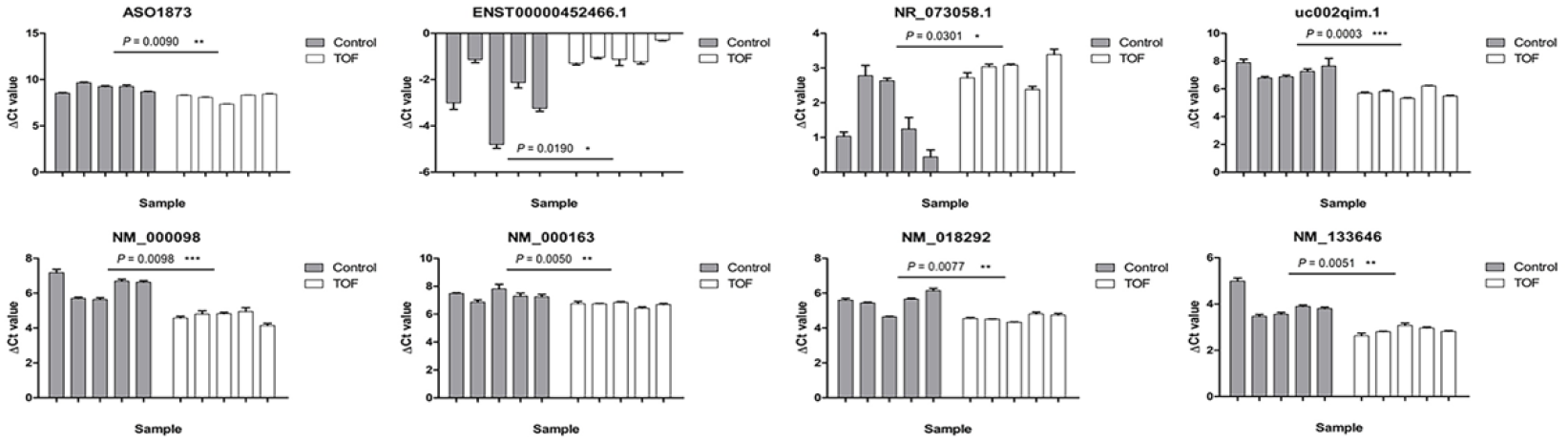
The qRT-PCR results of four lncRNAs and four mRNAs which are consistent with the chip results

## Discussion

TOF is the most common cyanotic congenital heart disease, accounting for approximately 0.2‰ *of* live births. Its pathological features include pulmonary artery stenosis, aortic straddle, ventricular septal defect, and right ventricular hypertrophy. The prognosis of the disease is closely related to the cause of the disease (whether there is genetic change). Some TOF cases are accompanied by the change in detectable genetic materials, including common genetic syndrome and changes in CNVs, SNPs, and Small InDels (Liu *et al.* 2016; Liu *et al.* 2017; Wang *et al.* 2017). The common genetic syndrome includes trisomy 21 syndrome, trisomy 18 syndrome, DiGeorge syndrome (more than 80% of cases have 22q11 microdeletion), and jaw–heart–face syndrome(Hou *et al.* 2019; Souto Filho *et al.* 2019). Patients with TOF accompanied by genetic material changes have different degrees of growth retardation, mental retardation, immune dysfunction, and malformation. After the follow-up, the prognosis of TOF patients with gene mutation is poorer than that of patients with TOF alone. At present, chromosome abnormalities, gene mutation, and other genetic mechanisms have been widely used in the diagnosis and treatment of TOF. For most patients with TOF, the condition involves changes in the heart and is not accompanied by detectable chromosomal abnormalities.

Cardiac development is a process of polygenic regulation. The genetic and epigenetic mechanisms such as nucleic acid modification and ncRNA also play an important role in its occurrence(Serra-Juhe *et al.* 2015; Grunert *et al.* 2016; Muntean *et al.* 2017). However, the role of genetic and epigenetic modifications in congenital heart disease should be studied in depth. The role of ncRNA in heart development has attracted increasing attention. Evidence shows that ncRNA can take part in the regulation of heart development-related pathways and lead to the occurrence of congenital heart disease(Wang *et al.* 2016; Turton *et al.* 2019; Wu *et al.* 2019). The studies on miRNA in the heart were carried out earlier and there were more studies on it. miRNA-1275, miRNA-27b, miRNA-421, miRNA-1201, and miRNA-122 may be associated with TOF; miRNA-27b may inhibit Mef2c, which affects myocardium development; miRNA-145 may regulate cell apoptosis and mitochondrial function by affecting the expression of FXN gene, thus leading to the occurrence of congenital heart disease. Studies on LncRNA have proved that lncRNAs exist in all stages of embryonic heart development. A very small amount of lncRNAs may be directly related to congenital heart disease. However, whole genome-based studies on lncRNAs, especially on intron or intergenic lncRNAs, are lacking. Given the diversity in the categories and functions of lncRNAs, the reference significance of study results on different lncRNAs is not high. In addition, given the lack of conservation among species, the expression level is generally low. There is no uniform name for lncRNA characteristics in the world, so there is difficulty in understanding its real meaning and function from the name. Given the absence of an lncRNA database, studying lncRNAs is difficult. Moreover, the role of lncRNAs in heart development should be elucidated. With progress in this field, many circRNAs have been identified to be distributed in many organisms; an evolutionarily conserved endogenous ncRNA was found to play a crucial part in the regulation of gene expression and participate in the occurrence and development of many diseases and embryonic development(Zhao *et al.* 2016; Liu *et al.* 2018). Studies on the role of circRNAs in the regulation of congenital heart disease are few. Thus, the role of circRNAs in congenital heart disease should be investigated. Thus far, the whole genome transcription of TOF has not been reported yet. In the present work, RNA was detected in the RVOT among patients with TOF. The differential lncRNA, miRNA, circRNA, and mRNA profiles of TOF are first reported comprehensively to have a basis for further exploring the etiology and individualized treatment of TOF.

In this work, 3228 lncRNAs which are differentially expressed, 4295 mRNAs which are differentially expressed, 118 miRNAs which are differentially expressed, and 18,016 circRNAs which are differentially expressed were found. Clustering analysis and principal component analysis were used to evaluate the reliability of data. The results show that in-group samples have a good correlation and consistency; between-group samples are significantly different and can be clearly grouped with strong data reliability. Next, target gene prediction, GO, and KEGG pathway enrichment analyses were conducted for differential locus. Five pairs of lncRNA–mRNAs were selected and verified via qPCR technology. Finally, four pairs of lncRNA–mRNA, namely, ASO1873-QRSL1, ENST00000452466.1-CPT2, NR_073058.1–GHR, and uc002qim.1-ZAK, were identified. Given the diversity in the categories and functions of lncRNA, the reference significance among different lncRNA study results is low. Candidate lncRNAs were identified on the basis of mRNA. The related literature shows that ZAK in four candidate genes is most likely related to TOF. ZAK is a mitogen-activated protein kinase kinase kinase (MAPKKK) which activates the stress-activated protein kinase/c-jun N-terminal kinase pathway and NF-κB. Twenty-seven different tissue samples from 95 individuals were tested via RNA-seq. The expression of ZAK in the heart is the highest. This gene, as a member of the MAPKKK family of signal transduction molecules, encodes a protein with an N-terminal kinase catalytic domain, which are followed by a leucine zipper motif and a sterile-alpha motif. This magnesium-binding protein, which forms homodimers, is located in the cytoplasm. The protein mediates gamma radiation signaling leading to cell cycle arrest, and the activity of which plays a role in cell cycle checkpoint regulation in cells. The protein also has pro-apoptotic activity(Jandhyala *et al.* 2008). ZAK can directly or indirectly affect the growth and apoptosis of cardiomyocytes. In 2004, Huang et al. reported that ZAK’s expression of a wild-type form induces the characteristic hypertrophic growth features, which include increased cell size, elevated atrial natriuretic factor (ANF) expression, and increased actin fiber organization(Huang *et al.* 2004a). In 2004, Huang et al. reported that ZAK’s expression of a dominant-negative form inhibited the characteristic TGF-β-induced features of cardiac hypertrophy, containing increased cell size, elevated expression of ANF, and increased organization of actin fibers(Huang *et al.* 2004b). In 2016, Fu et al. reported that ZAKα signaling activation is crucial for cardiac hypertrophy, and their findings revealed the inherent regulatory role of ZAKβ in antagonizing ZAKα and subsequently downregulating the cardiac hypertrophy and apoptosis induced by ZAKα (Fu *et al.* 2016). After selecting ZAK, lncRNA uc002qim.1 was selected and may affect ZAK expression on the basis of the target gene prediction result. The selected miRNA was consistent and interacted with target mRNA in the expression trend. Thus, hsa-miR-130b-3p, hsa-miR-372-3p, and hsa-miR-758-3p were miRNAs that may get involved in the lncRNA–mRNA interaction. The miRNA’s seed sequences should be compared with lncRNA to further understand their interaction relationship. Through comparison of seed sequences, lncRNA uc002qim.1 has three potential binding loci with hsa-miR-130b-3p, two potential binding loci with hsa-miR-372-3p, and one potential binding locus with hsa-miR-758-3p. Therefore, hsa-miR-130b-3p may likely play a regulatory role in lncRNA uc002qim.1-mRNA ZAK interaction. The relationship between hsa-miR-130b-3p and congenital heart disease has not been established before. However, hsa-miR-130b-3p has been widely studied. Some studies have shown that hsa-miR-130b-3p can regulate the differentiation of keratinocytes and play a significant part in the skin disease formation(Li *et al.* 2017). miR-130b-3p also acts a pivotal part in the invasion and differentiation of tumor cells. It can inhibit the breast cancer cells’ invasion and migration by targeting Notch ligand Delta like-1. miR-130b-3p can negatively regulate PTEN through PI3K and integrin β1 signaling pathway, thereby playing a part in bladder cancer’s occurrence and development(Yu *et al.* 2015; Shui *et al.* 2017). Thus, the interaction among uc002qim.1, ZAK, and miR-130b-3p should be studied.

In this work, whole genome chip technology screening found that circRNA has the largest data volume. A total of 18,016 circRNAs, which are differentially expressed, including 9404 upregulated and 8612 downregulated circRNAs, were screened. Target miRNA interaction analysis was conducted for the first eight circRNAs, and the interactive network was constructed for further studies. We could not observe the interaction of circRNA with uc002qim.1, ZAK, and miR-130b-3p. cirRNA may have an effect on heart development through other unknown pathways.

In summary, the patterns of expression of lncRNAs, mRNAs, miRNAs, and circRNAs in the RVOT of patients with TOF were studied using Agilent human lncRNA+mRNA Array v4.0, Agilent human miRNA Array V21.0, and Agilent human circRNA Array v2.0. In this paper, the full transcriptome data of RVOT myocardial tissue among patients with TOF were systematically reported for the first time, and differentiation, GO, KEGG pathway enrichment, and target gene prediction analyses were conducted to obtain these data. Finally, the optimal RNA sequencing information was provided. As such, the function of these ncRNAs, their molecular regulatory mechanisms, and their pathological mechanism in the occurrence and development of simple TOF can be analyzed. Given sample size limitations, this study should be further strengthened. The function and regulatory mechanism of ncRNAs should be further studied.

## Author Contributions

In this study, HDW and LL conceived the concept of the work and designed the study. CYC, YNL, YYL, collected the data and performed the statistical analyses. TBF and BTP collected the sample. HDW and LL revised and finalized the manuscript. All authors read and approved the final manuscript.

## Acknowledgments

We are grateful to Hao Tang and Yanze Li for offering the technical support.

